# CRISPR-MIP replaces PCR and reveals GC and oversampling bias in pooled CRISPR screens

**DOI:** 10.1101/2024.03.28.587082

**Authors:** Martin Selinger, Iryna Yakovenko, Iqra Nazir, Johan Henriksson

**Affiliations:** Department of Molecular Biology, Umeå University, Sweden; Laboratory for Molecular Infection Medicine (MIMS), Sweden

## Abstract

Pooled CRISPR screening is a powerful tool for finding the most important genes related to a biological process of interest. The quality of the generated gene list is however influenced by a range of technical parameters, such as CRISPR (single guide) sgRNA target efficiency, and further innovations are still called for. One open problem is the precise estimation of sgRNA abundances, as required for the statistical analysis. We do so using molecular inversion probes (MIPs) combined with the use of unique molecular identifiers (UMIs), thus enabling deduplication and absolute counting of cells. We show that this is a viable approach that eliminates sequencing depth bias. Furthermore, we find that GC% bias affects PCR, calling for a reanalysis of published CRISPR screen data and sgRNA efficiency estimates. We propose our method as a new gold standard for sgRNA quantification, especially for genes that are not top ranked but still of broad interest.

## Introduction

Pooled CRISPR screens are a rapid and comparably cheap method to evaluate which genes are involved in a particular biological process. The typical process is illustrated in Figure 1A; using pooled cloning, typically lentiviruses are produced that each are capable of expressing on single guide (sg)RNA. After packaging, cells of interest are transduced such that each cell has at most one virus integrated. Combined with Cas9 (which can be delivered in several ways), this causes CRISPR-based knock out of a random target gene. Cells are selected based on a phenotype of interest, which may be altered due to the knockout. To find which sgRNA caused the alteration, genomic DNA (gDNA) is extracted, and the integrated sgRNA sequence PCR-amplified. High-throughput sequencing (typically Illumina-based) is then used to count the sgRNA abundances for the cells, as well as a reference sample.

**Figure 1.**
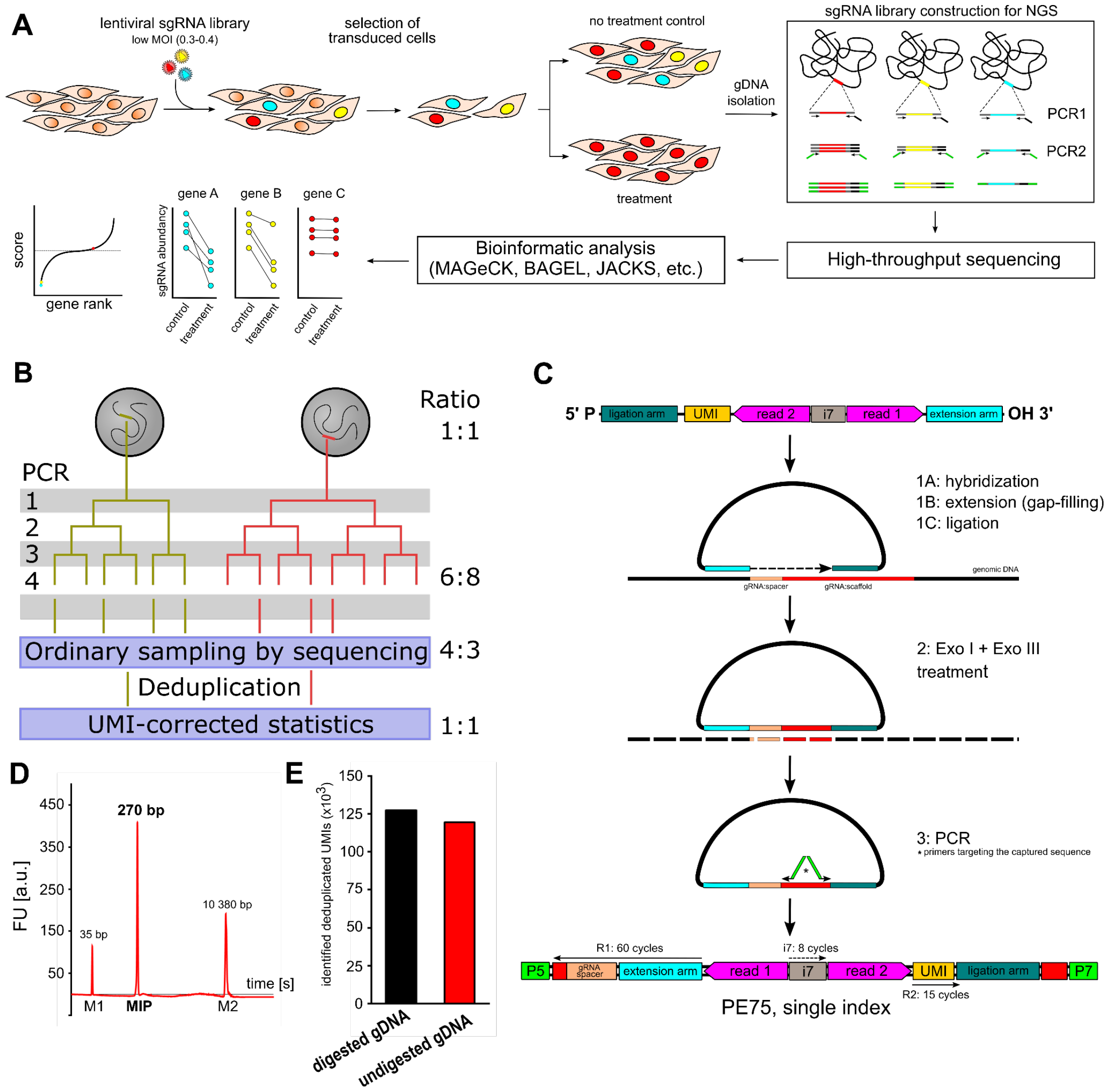
CRISPR sgRNA can be amplified by molecular inversion probes. **(a)** Overview of a typical pooled CRISPR screen. **(b)** Overview of the steps in a PCR amplification and sequencing process, showing how the final and intermediate ratios differ from the true sgRNA count ratios. Because the sequencer samples multiple molecules from the same source (yellow *vs* red), these are not independent events in a probabilistic sense. This makes statistical modeling difficult, as it normally assumes i.i.d (independent identically distributed) variables. This can be corrected using UMIs but these cannot be added to a regular genomic PCR protocol. **(c)** Overview of the CRISPR-MIP protocol. **(d)** A bioanalyzer trace of the final product shows clean and specific amplification (1 sgRNA CRISPR screen model). **(e)** Number of unique UMIs identified after deduplication (umi-tools) in case of HindIII/BamHI-digested gDNA and undigested gDNA; input amount of gDNA was equivalent to 150 000 cells.

In practice, the pooled screening approach can be difficult to perform, and the success relies on a large number of technical details. These include potential difficulty of transduction and efficiency of genome editing, which we have especially noted for primary cells^1^. Overall it may be difficult to produce a large number of edited cells. Having a finite number of integrated sgRNAs to quantify makes reliable statistics challenging. The issue has partially been circumvented by, e.g., improved computational approaches^2^ and lineage tracing viruses^3,4^. The latter are especially promising but have been reported to be difficult to use in practice. This mainly stems from the requirement to generate ultra complex viral libraries, where the complexity can be lost at any step in the cloning and transfection process. Even in the ideal scenario, the lineage barcodes cannot be guaranteed to be unique, as plasmids are typically grown after the final cloning step, before virus production.

We have approached the problem from an orthogonal angle, which can be combined with the previous methods. We hypothesize that the PCR amplification of integrated sgRNAs, the last step in a CRISPR screen, may negatively affect the statistical robustness of CRISPR screens. For example, only relative abundances of sgRNA PCR copies are extracted, but these need not perfectly reflect cell counts (Figure 1B). Possibly worse is that the number of sequencing reads, after PCR, is much greater than the number of cells. Thus the same initial sgRNA is sampled through multiple sgRNA PCR copies, making the reads statistically dependent events. This violates the Poisson sampling assumption for the sgRNA read counts, which is the basis of most sequencing-type statistics. The negative binomial approximation is commonly used generalization to better handle the relationship between the count mean and the dispersion (variance), such as to calculate p-values and fold changes for RNA-seq differential expression^5^. It is however not obvious that this generalization is the best model for pooled CRISPR screen data.

To circumvent this problem, we borrow the concept of UMIs (unique molecular identifiers) which are especially ubiquitous in the single-cell field. UMIs allow deduplication of reads of the same original molecule, thereby removing PCR bias and improving statistics^6^. Since UMIs cannot be used for a basic PCR as done today, we instead implement these using MIP (molecular inversion probe)-based amplification. MIPs represent a highly efficient and specific technique for targeted genomic enrichment and sequencing. Initially conceptualized for the detection of single nucleotide polymorphisms, MIPs have since evolved to facilitate a wide array of genomic analyses, including large-scale genotyping, copy number variation detection, and targeted sequencing of specific genomic regions. These probes uniquely enable the selective amplification and sequencing of desired DNA fragments, leveraging a circularization-based capture mechanism that ensures high specificity and sensitivity.

In this paper we present a novel MIP-based approach, which to our knowledge has never been used for this application. We provide a novel design that eliminates the common problem of huge background noise. Our protocol is simple and is a drop-in replacement of PCR, and it can be used in conjunction with previous protocols for even further improved precision. We thus believe our approach will be of broad utility to the CRISPR screening field.

**Supplemental Figure 1.**
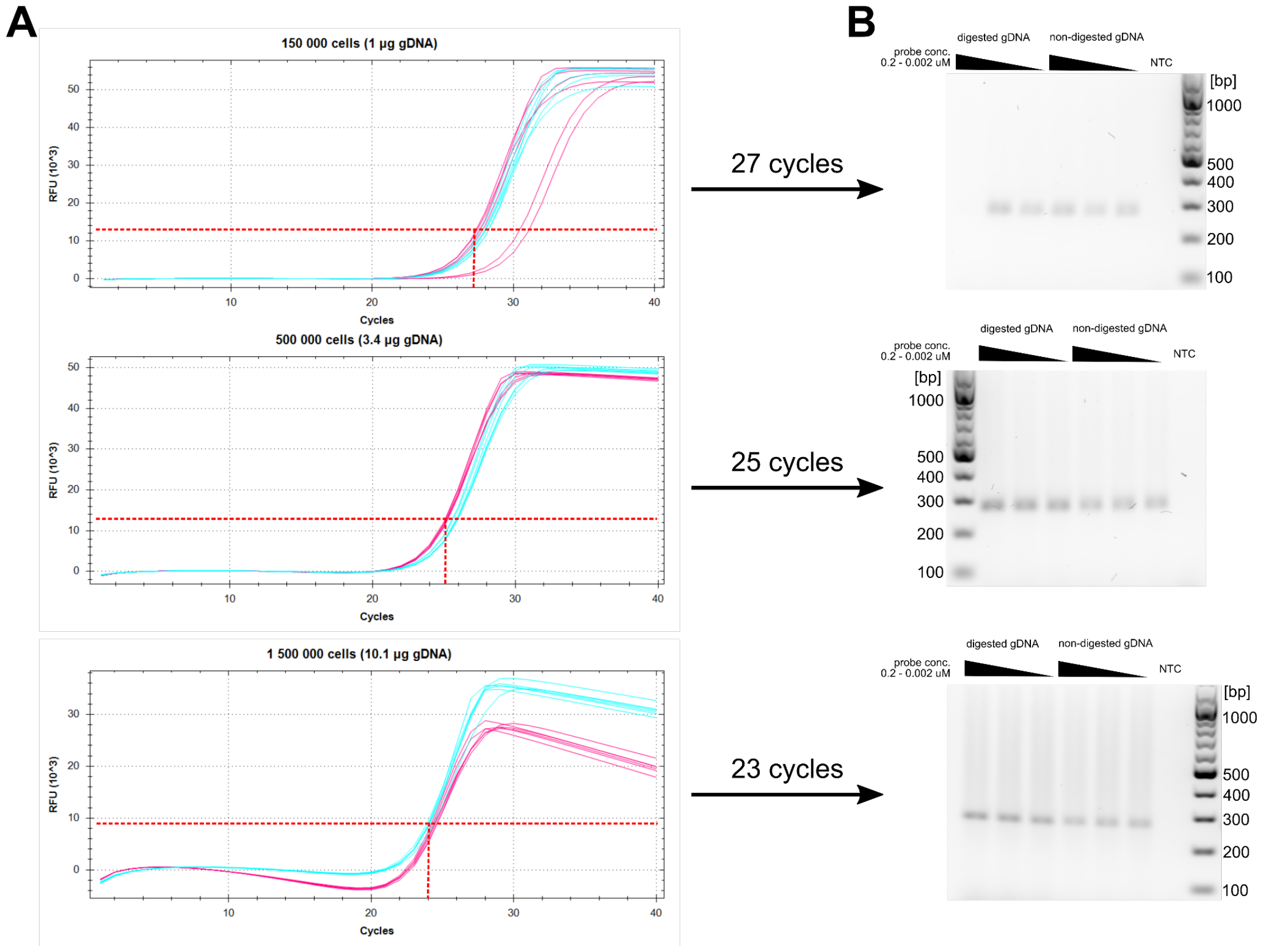
Optimization of CRISPR-MIP protocol. **(a)** qPCR amplification curves for CRISPR-MIP protocol using different input amounts for HindIII/BamHI-digested gDNA (pink) and undigested gDNA (turquoise). **(b)** Analysis of fragment amplification based on qPCR.

### CRISPR sgRNAs can be quantified by molecular inversion probes

Our CRISPR-MIP design is shown in Figure 1C and is inspired by previous MIP designs^7–9^. We make use of the fact that most sgRNA libraries are based on U6 promoter driven expression, and that sgRNAs have a common scaffold sequence. Thus a single MIP can target all sgRNAs within the library. To capture the sgRNA target sequence, we perform a fill-in using Phusion polymerase, followed by high-temperature (60°C) ligation to circularize the construct using a thermostable ligase. Because the gDNA sampling can only happen once, unlike PCR, this enables the addition of a UMI in the MIP backbone. After cleanup, circularization and digestion of the corresponding single DNA strand, the MIP is then amplified by regular PCR, which also adds P5/P7 adapters for Illumina sequencing. The probe backbone also contains an i7 index for sample multiplexing. To overcome any background caused by circularized “empty” probes, both PCR primers are targeted against part of the captured sequence itself (Figure 1C) instead of the MIP backbone.

A difference to previous designs is that we took particular care in the design to enable sample multiplexing with other types of libraries, i.e., it can be pooled with other routine Nextera or Truseq-type libraries (i.e. most regular Illumina libraries). This meant also that we could sequence our libraries with the PCR-prepared libraries for comparison. Furthermore, the probe can be covered by 60/15 bp paired end, making it fit with the cheapest Illumina 75 cycle kits. Together, this makes our CRISPR-MIP design maximally cost effective on modern flow cells, which require large numbers of samples to be multiplexed.

Initial optimization was performed on a one-sgRNA CRISPR screen model utilizing the lentiCRISPRv2 system in pancreatic cancer cell line MiaPaCa-2 (see methods). While a 1:1 ratio of gDNA:MIP is theoretically optimal, frequently a much larger amount of MIPs are required^7–9^. After optimization, 0.2 pmoles of probe per reaction were proved to be optimal for 0.5 - 1.0 × 10^6^ cells (gDNA:MIP ratio 1:120×10^3^ - 1:240×10^3^; Figure S1). The final protocol proved to be highly specific with no background (i.e. absence of any off-products of the wrong size; Figure 1D, Figure S1B). We also speculated that digesting the gDNA would open up the chromatin and thus improve MIP annealing efficiency; however, the UMI-deduplicated data comparison documented only a minor difference (Figure 1E). In addition, the digestion step was further offset by the need for cleanup and buffer exchange.

**Supplemental Figure 2.**
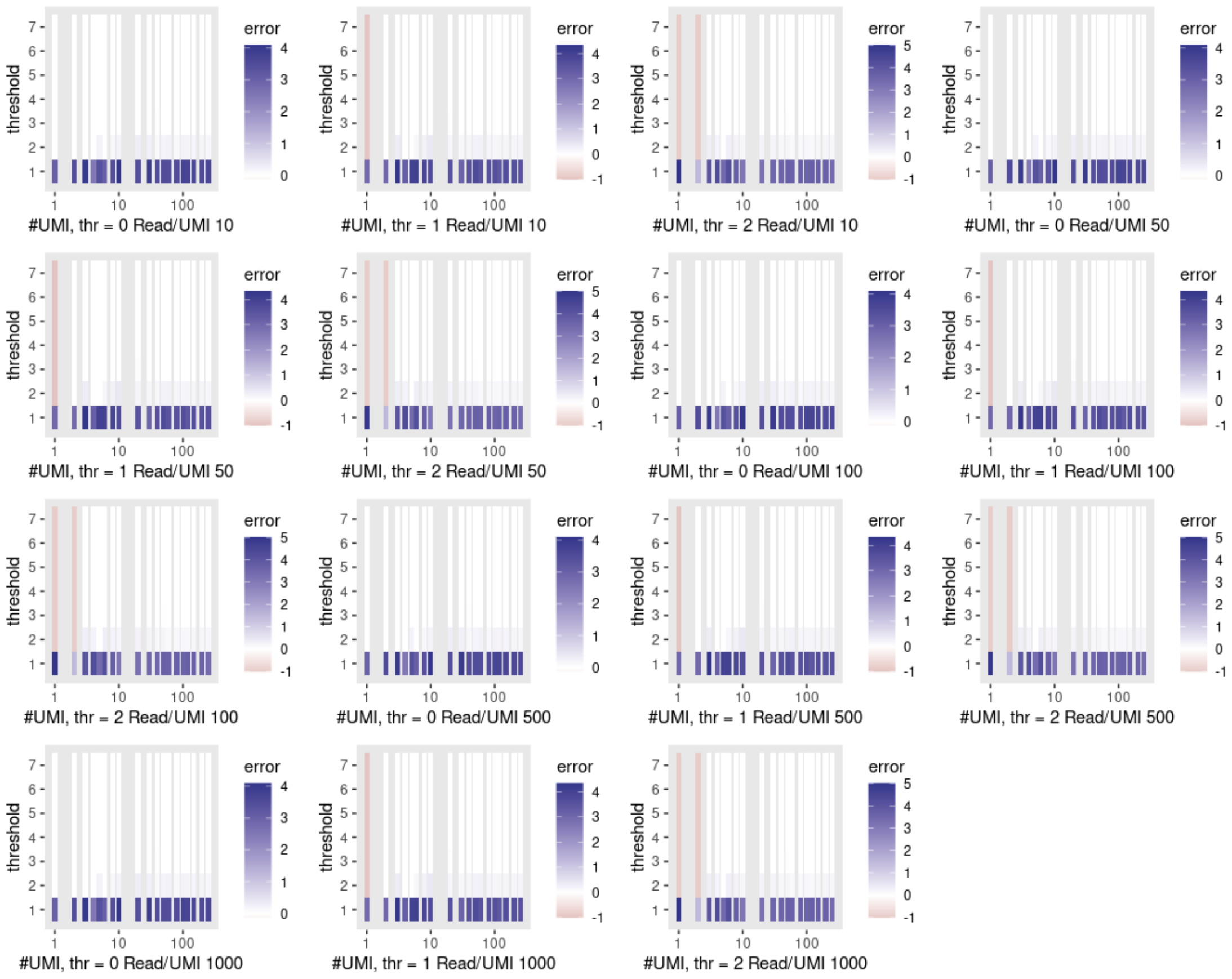
Optimization of UMI deduplication algorithm. Reconstruction error after UMI deduplication, for different number of UMIs (x-axis), UMI-tools threshold (y-axis), number of UMIs in the space, minimum threshold (thr) for keeping a group of UMIs, and number of sequencing reads per UMI. The error is defined as (DetectedUMIs - ActualUMI) / ActualUMIs, and is in a range [-1,inf).

**Supplemental Figure 3.**
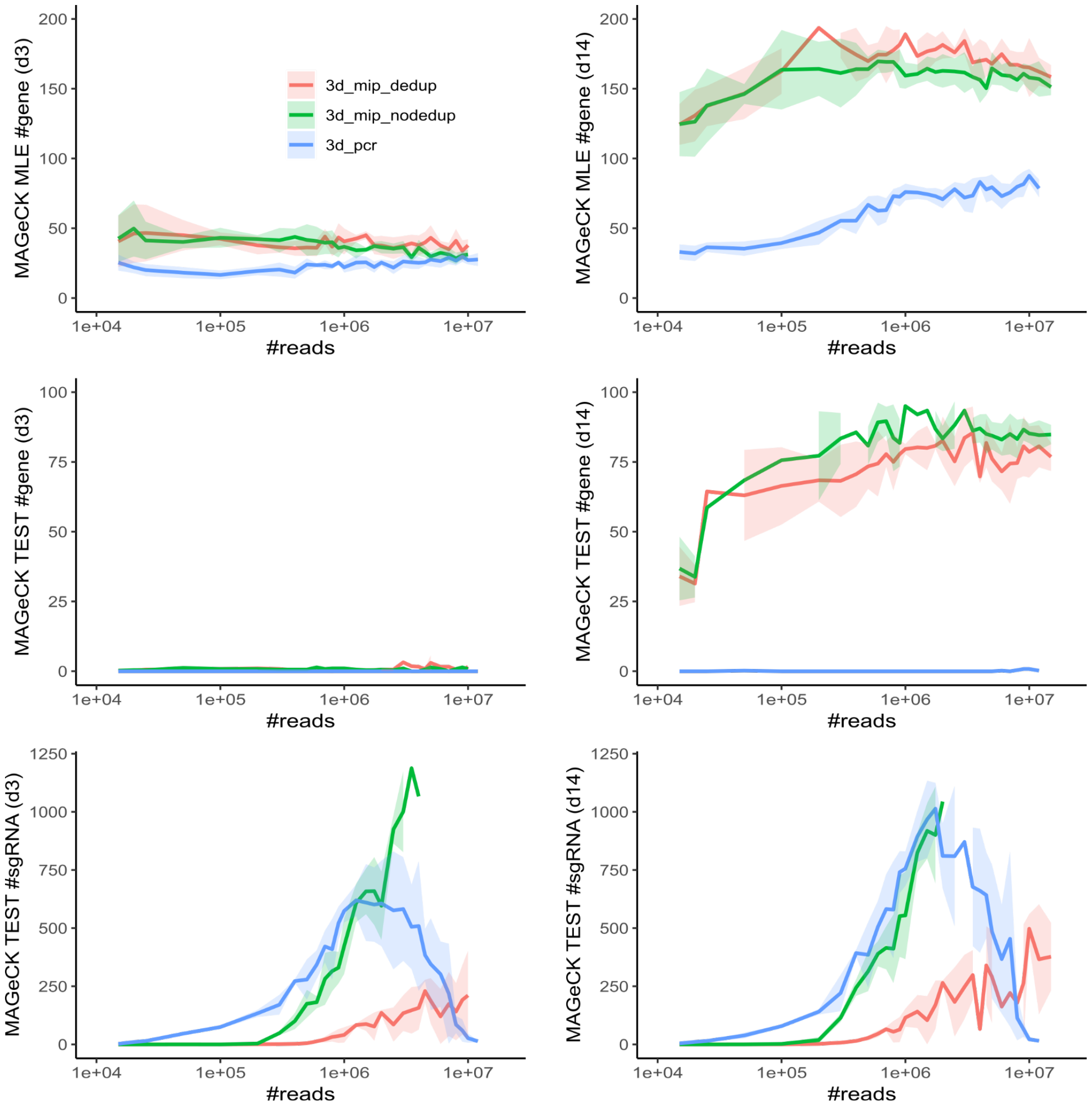
Full statistics for day 3 and 14 in the dropout screen. Comparisons of CRISPR-MIP and PCR, using both MAGeCK test and MLE modes of estimation. Number of significant genes (FDR<0.05) and sgRNAs (FDR<1e-5).

### CRISPR-MIP is resilient to sequencing depth variation and saturation

To test our new method in a real scenario, we conducted a dropout screen in MiaPaCa-2 cells using the Brunello kinome library targeting 763 human kinase genes^10^. To assess the performance of CRISPR-MIP relative to the conventional PCR protocol, sgRNA libraries from each sample were prepared by both techniques (Figure 2A). We then implemented a custom counter for CRISPR-MIP that takes care of UMI-based deduplication, making it ready for regular downstream analysis such as MAGeCK hit calling (Figure 2B). The deduplication is based on UMI-tools^11^. To tune the parameters, we investigated in particular how sequencing errors affect deduplication. We found 3.5% mismatches/bp in one typical run, and show the empirical distribution UMIs of one sgRNA *vs* theoretical estimates (Figure 2C). Most associated reads are at most 2-3 bp different due to sequencing errors, and are still distinct from a second UMI. Because UMI-tools use further heuristics for corrections, we also performed a simulation to account for UMI space saturation, and the possibility of removing reads that cannot be placed in any particular group (Figure 2D, S2). We find that deduplication can be near perfect for threshold = 2 for up to 200 UMIs/sgRNA, and 10-1000 sequencing reads per UMI. We thus recommend this setting, and ideally 10x more reads than cells used in the screen (e.g. 1M cells ⇒ 10M reads).

**Figure 2.**
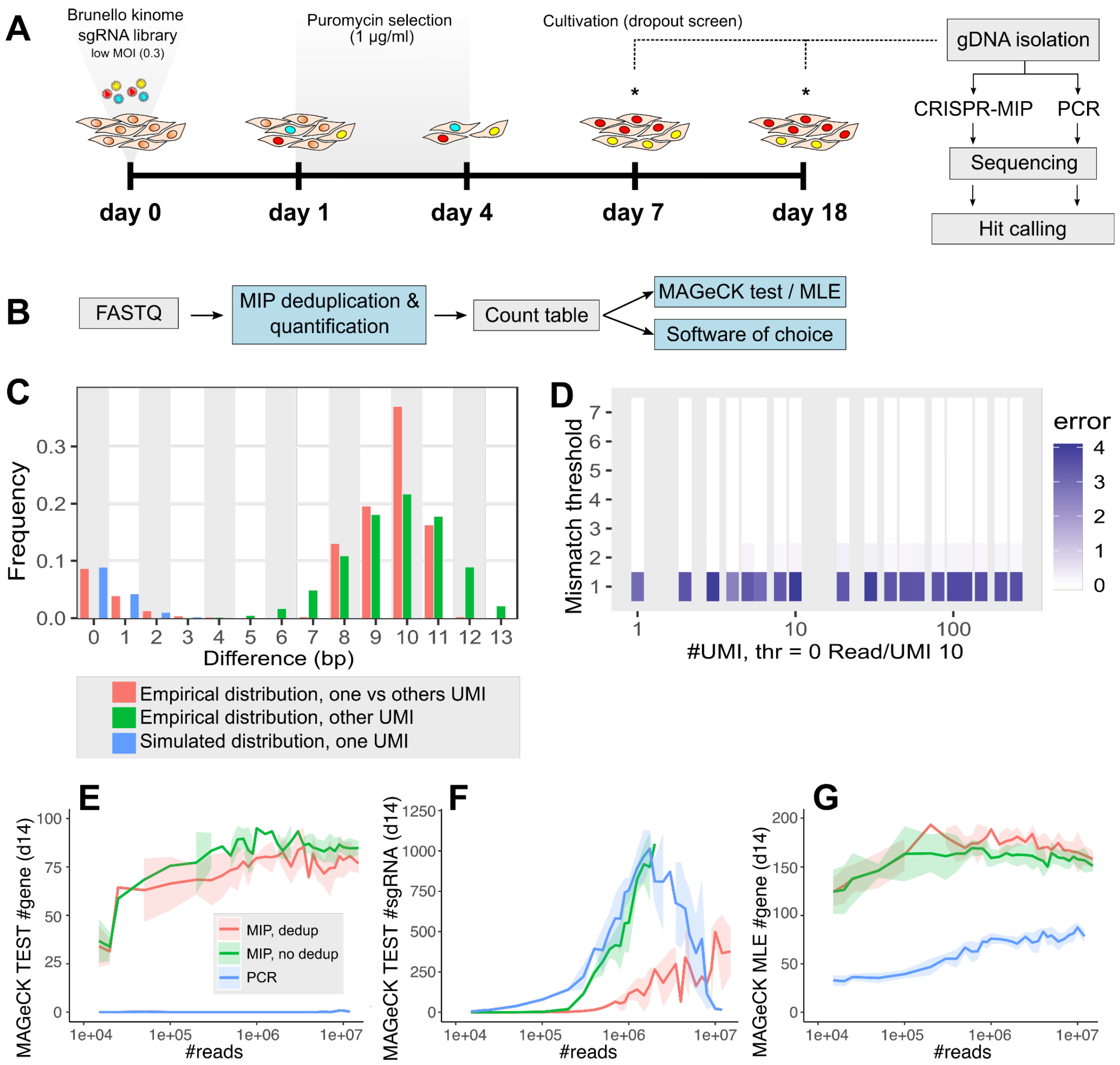
CRISPR-MIP is resilient to sequencing depth variation and saturation. **(a)** Experimental setup for our dropout screen. Each sample was prepared in biological triplicates. **(b)** Our deduplication pipeline generates a count table that can be analyzed using common software for pooled CRISPR screen statistical analysis. **(c)** Analysis of distance between UMIs (number of bp different): Empirical distribution of distance from most abundant UMI to all other reads (red), fitted distribution to read abundances for one abundant sgRNA (green) and theoretical distribution from one single UMI, to all other possible UMIs, ignoring sequencing errors (red). **(d)** Recovery of initial molecules as a function of reads/cell, after UMI correction. The number of UMIs (x-axis) and UMI-tools threshold parameter (y-axis). All groups of UMIs found by UMI-tools are retained. The error is defined as (DetectedUMIs - ActualUMIs) / ActualUMIs, and is in a range [-1,inf). **(e)** Number of significant sgRNAs (FDR < 1e-5) for PCR, and MIPs with and without UMI-correction. PCR-based analysis has a particularly odd trend *vs* the depth, making it unclear how to interpret the statistical output. **(f)** Number of significant genes (FDR < 0.05) for PCR, and MIPs with and without UMI-correction.

With our optimized deduplication algorithm, we next compared the statistical output to PCR as a function of sequencing depth. We focused on day 14 as it had more significant genes than day 3, but the conclusions are the same (Figure S3). We note that PCR results in many more sgRNAs being significant (Figure 2E), but oddly only for intermediate numbers of reads. This makes it unclear how to interpret results from PCR. MIP without deduplication also results in more significant genes, but the number is at least monotonously increasing with sequencing depth. Both of these trends suggest inflated statistics when the origin of reads is not resolved, in accordance with our hypothesis. This is further reinforced when comparing the low number of significant genes (Figure 2F), which is much lower in PCR, contradictory to many significant sgRNAs. The alternative MAGeCK MLE model handles this situation somewhat better by consistently finding more hits at higher sequencing depth also for PCR, but finds fewer significant genes (Figure 2G).

To summarize, we show that deduplicated MIPs have superior robustness to sequencing depth. CRISPR-MIP especially appears to give increasingly better results, for larger sequencing depth, until the complexity of the library becomes the bottleneck - an expected basic property which unfortunately the PCR approach fails. Thus we have exposed a serious flaw in the PCR protocol, and provide CRISPR-MIP as a solution.

### CRISPR-MIP finds genes which are not retained by PCR

Next we analyzed the functional output from the kinome dropout screen. For this, we analyzed median-normalized, full-scale sequencing data from CRISPR-MIP and PCR libraries using both the MAGeCK test and MLE algorithms^12,13^. Prior to the analysis, UMI deduplication was performed in the case of CRISPR-MIP to enable absolute cell counting and improved statistics. In accordance with the subsampling analyses, CRISPR-MIP demonstrated a higher sensitivity in detecting statistically significant hits (FDR < 0.05) compared to PCR, with CRISPR-MIP identifying 63 and 68 genes for MAGeCK MLE and test, respectively, versus 20 and 2 genes for PCR (Table S1). Despite the large differences, genes such as CDK1, PLK1, and CDC7 are consistently ranked as top significant hits (Figure 3A). The methods also agree on the most significant GO terms (Figure 3B) which are all related to cell cycle and growth, as expected. Furthermore, the RRA/beta score ranks are also highly correlated (correlation coefficients of 0.83 and 0.79) (Figure 3C). This raises the questions as for why so few significant genes are detected for PCR.

**Figure 3.**
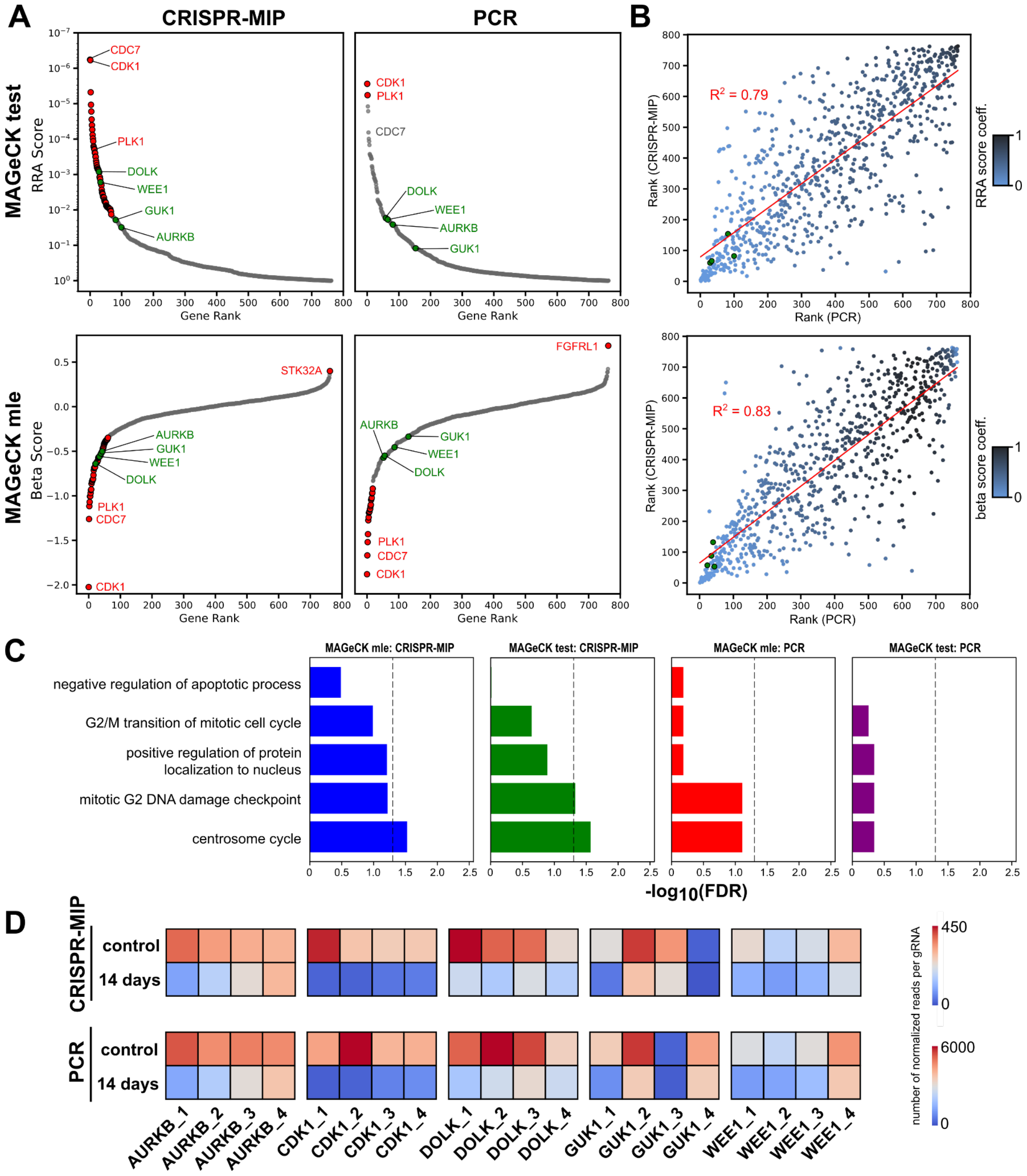
CRISPR-MIP detects valid hits not retained by PCR. **(a)** Comparison of statistically significant hits (red; FDR < 0.05) between CRISPR-MIP and PCR using MAGeCK test and MLE algorithms. CDK1, CDC7, and PLK1 represent top-ranked genes with high statistical significance among all analyses, whereas genes AURKB, DOLK, GUK1, and WEE1 (green) represent a group with variable significance. **(b)** Correlation of ranks between CRISPR-MIP and PCR (genes AURKB, DOLK, GUK1, and WEE1 highlighted in green). The beta/RRA score coefficient = fdr_CRISPR-MIP_ * fdr_PCR_. **(c)** Top GO:BP terms for genes listed as significant hits for CRISPR-MIP and PCR. Dotted lines represent a significance threshold of FDR < 0.05. **(d)** Heatmap illustrating normalized read numbers for each of the 4 sgRNAs present in Brunello kinome library in case of genes with consistent top hit (CDK1) and variable hits (AURKB, DOLK, GUK1, WEE1).

To better understand the differences between CRISPR-MIP and PCR, we investigated some genes which agreed less in significance between the two methods; These include AURKB, WEE1, and GUK1, scored as highly essential in DepMap^14^. We also found that MIP pinpoints DOLK, which is lowly scored in DepMap^14^. Detailed analysis of individual sgRNA abundances show a high level of agreement between the two protocols (Figure 3D). Together with the low significance for PCR, this suggests that the low number of significant genes from PCR might primarily be due to the estimation of the variance rather than the effect size.

To summarize, CRISPR-MIP gives results not too different from PCR for the top hits. The main improvement is in improved statistical estimation. However, the improved precision might also lead to further novel findings, as exemplified by the gene DOLK. Thus to uncover genes beyond the most obvious, such as CDK1 (the canonical cell cycle and growth control gene), improved detection methods such as CRISPR-MIP will be increasingly beneficial.

### CRISPR-MIP avoids widespread PCR-induced variance and systematic bias

A key question is *why* the MIPs find more significant hits than PCR. A closer look at the abundances of sgRNAs show that they are rather similar for the genes we investigated, PCR *vs* CRISPR-MIP (Figure 4a). However, comparing MIP against PCR globally, there are some particularly extreme deviations (Figure 4b). Detection of CRISPR screen hits is similar to differential expression in that a value akin to a z-score must be calculated, to in turn generate a p-value. The z-score is of the form *z = μ/σ*, and thus the mean effect size and the dispersion must be estimated. Since the effect size does not seem to differ between the library preparation methods, at least for some genes that we show are true hits, this leaves the dispersion estimation as the potential source of the problem. The seeming larger variation of abundances for PCR is thus affecting the scoring of all genes globally.

**Figure 4.**
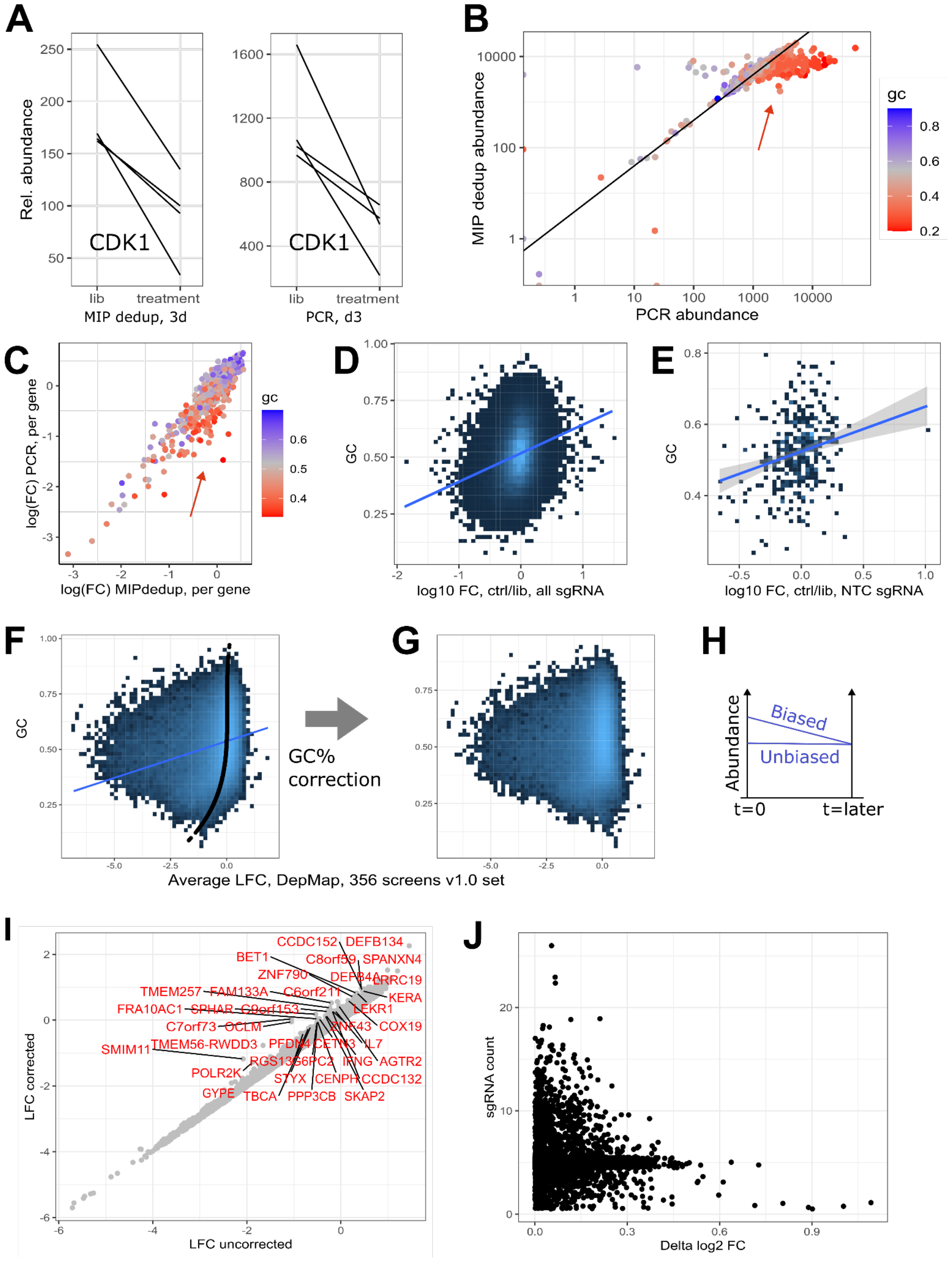
CRISPR-MIP avoids widespread PCR-induced variance and systematic bias. **(a)** Library sgRNA abundances of one gene, showing the similar effect independent of library preparation method. **(b)** Comparison of sgRNA abundances according to MIP and PCR amplification, library *vs* day 3 (d3). **(c)** Comparison of gene fold changes according to MIP and PCR amplification, library *vs* d3. **(d)** Abundances of all sgRNAs from Sintov2022^17^ vs GC%. **(e)** Abundances of non-target control (NTC) sgRNAs from Sintov2022^17^ *vs* GC%. **(f)** DepMap^14^ pooled CRISPR screens average fold changes vs GC%; fitted mean in black; linear model in blue. **(g)** DepMap, after GC% correction. **(h)** Overestimation of sgRNA abundance in the library will lead to artificially essential genes in a drop-out screen. **(i)** Fold changes of genes, before *vs* after GC correction. **(j)** Difference in fold change *vs* number of sgRNAs per gene, showing that differences can be large also for 10+ sgRNA/gene.

We looked for reasons that could explain the larger variance of PCR, and discovered that GC% has a high impact on abundance (Figure 4b). Those with low GC% are most highly abundant. We find this somewhat surprising given that the amplicons only differ in 20bp. However, the effect of GC% on PCR efficiency is well-known, and complex. In the context of metagenomics, sequences with high GC% underrepresented, but those with low GC% are even more so^18^. Interestingly, a strong bias is not present for PCR on gDNA (Figure 4c). If naked plasmid DNA is more accessible than gDNA, which can be due to differences in coiling, or remnant proteins, then the melting temperature can be the rate limiting factor for amplification. We reason that low GC%, and thus low melting temperature, has an effect in the first cycles of PCR. If, for example, it takes 3 extra cycles to successfully copy one input molecule, then it artificially lags 2^3^=8 times behind in abundance estimation. This or a similar hypothesis would explain the large technical variation of PCR. Since MIPs can only amplify the gDNA once, during the circularization step, it is either all-or-nothing. The UMI-deduplication also removes any PCR-bias in subsequent steps.

To validate our findings we also investigated public data. We found a larger lentiCRISPRv2-based screen on hESCs that includes both library PCR and an initial day 10+ time point (Sintov2022)^17^. Comparing this much larger number of sgRNAs with GC%, we see a strong similar trend (Figure 4d). To rule out that the trend is due to differential gene editing efficiency of sgRNAs, we also looked at the subset of 1000 non-targeting controls (NTCs), and still see a similar trend (Figure 4e).

Sintov2022 compared gDNA at a late time point, to that of an earlier time point, likely avoiding plasmid PCR-induced biases, we became concerned if there are screens that instead do use plasmid PCR as the reference. Scanning the literature, we noticed it to be the case for a large dropout screen dataset, covering 324 human cancer cell lines with the aim of finding core and cancer specific essential genes (DepMap)^14^. Again, we indeed find a GC-skew in sgRNA fold change library *vs* average dropout across all screens (Figure 4F, thus corresponding to “core” essential genes). We reason that plasmid PCR bias should lead to some genes having artificially high essentiality, as abundances will be overestimated for the reference plasmid (Figure 4H). To estimate the effect not on individual sgRNAs, but genes, we fitted a simple model to undo the effect of the GC% bias (Figure 4G). The difference in average fold change, for each gene, is then shown in Fig 4i. Unsurprisingly, the most sensitive genes to GC-bias are those that are already hard to reliably estimate: Those having low fold-change, and/or few sgRNAs (Figure 4J). However, differences exist sometimes even for >10 sgRNAs. Thus depending on the application, care must always be taken in interpreting screens if plasmid PCR was used as a reference.

## Discussion

There are many technical details that must be considered to perform a high-quality pooled CRISPR screen. Here we have focused on, and successfully resolved, the statistical issue of not knowing the precise copy number of sgRNAs when using PCR for library preparation. Not accounting for this resulted in seemingly unreliable statistics, at least for the prototypical screen considered here (Figure 2E). Our protocol instead has an expected and attractive property: deeper sequencing yields better statistics, up to the amount of cells sampled in the screen. Our method does not compete with methods that count independently editing clones^3,4^, but is rather a complement to make the final cell counting more robust. It however has the benefit of being simple to implement, without the need for hypercomplex pooled libraries.

Inadvertently, we have also discovered and resolved another problem: sgRNAs with unbalanced GC% are misrepresented by PCR, which can lead to issues if comparing gDNA to plasmid. We have found one example study where this has affected the results. This does not affect top hits in the screen, but there are two cases when all genes are analyzed, and thus this becomes a concern: (1) Gene regulatory network reconstruction, where phenotypes of all genes are included to order them in the cascade; (2) Optimization of sgRNA design^19^. If the GC% bias were to be included in ML models, it could inappropriately be considered to affect the gene editing efficiency. Designing good sgRNAs is already difficult due to constraints, e.g., the first 5bp next to the PAM direct Cas9 targeting^20^, making it difficult to find unique sites. Furthermore, sgRNAs can be chosen to maximize microhomology-directed repair to improve knockout efficiency; or otherwise optimized for homology based insertions^21^. Our MIP protocol at least alleviates further constraints due to PCR bias.

CRISPR-MIP may further help study the most essential genes. One workaround for the PCR-on-plasmid issue is to instead perform PCR of gDNA from a “time 0” time point, as is recommended in various protocols^22^. However, key genes may by this time already have dropped out of the library. It is thus better to use the plasmid as the reference, using CRISPR-MIP to avoid GC% bias. In either case, we strongly discourage PCR on plasmids for anything but quality control of the cloning.

To some extent, we believe improved computational methods can account for GC% bias in existing screens. However, investigating public data, we frequently find the method descriptions to be subpar, and raw sgRNA counts missing. This makes retrospective statistical improvements impossible. We thus urge the community to improve their FAIR standards such that we all can benefit from improved screening data.

## Methods

### Cell culture

HEK293T cells were obtained from the American Tissue Culture Collection (ATCC). MiaPaCa-2 and HEK293T cells were grown in high glucose DMEM supplemented with 10% FBS, 1% GlutaMAX (200 mM L-alanyl-L-glutamine dipeptide in 0.85% NaCl), and 1% antibiotics-antimycotics (amphotericin B 0.25 μg/ml, penicillin G 100 units/ml, streptomycin 100 μg/ml), all purchased from ThermoFisher Scientific. Cells were cultured at 37°C in 5% CO_2_ atmosphere and confirmed to be negative for mycoplasma contamination prior to the experiments.

### CRISPR/Cas9 screens

#### Plasmids

Human Kinome CRISPR pooled library (Brunello) was a gift from John Doench & David Root (Addgene #75314)^10^. Backbone plasmid in the library is lentiCRISPRv2 (Addgene #52961), which contains a U6-sgRNA expression cassette coupled with Cas9 gene as well as puromycin resistance gene for the selection of transduced cells. For lentivirus production pCMV-VSV-G (envelope plasmid) and pCMV dR8.2 dvpr (packaging plasmid) were gifts from Bob Weinberg (Addgene # 8454 and #8455, respectively)^25^. For the initial screen with GFP-targeting sgRNA, lentiCRISPRv2-GFPg1 plasmid was used. The Brunello Kinome library was amplified according to the BROAD Institute Amplification of pDNA Libraries Protocol and their complexity was subsequently verified by NGS sequencing performed according to the BROAD Institute PCR of sgRNAs for Illumina Sequencing. Primers used are in Table S2. Sequence of GFP-targeting sgRNA was derived from LentiGuide-Puro-GFPg1 (BB09)^24^

#### Lentivirus production

Lentiviruses were produced in HEK293T cells using Lipofectamine LTX transfection reagent (Invitrogen #A12621) according to the manufacturer’s instructions. Briefly, 5×10^6^ cells per 100 mm culture dish were seeded one day prior to the transfection. Next day, culture medium was exchanged for 5.7 ml of fresh pre-warmed media. The transfection mix per culture dish was prepared by combining 1 ml of premix 1 (5.67 μg of plasmid library, 2.84 μg of pCMV-VSV-G envelope plasmid, 5.67 μg of pCMV dR8.2 dvpr packaging plasmid in OptiMEM medium, Gibco #31985062) and 1 ml of premix 2 (42.5 μl of Lipofectamine LTX, 957.5 μl of OptiMEM media; DNA:LTX ratio 1:3, w/v). Transfection mix was subsequently incubated for 30 minutes at room temperature and added to the cells. After 6 hours, medium was exchanged by 10 mL of fresh pre-warmed media supplemented with 1× ViralBoost Reagent (Alstem, #VB100).

At 3 days post transfection, clarified medium supernatant (1600×g, 10 minutes, 4°C) was filtered through a 0.45 μm syringe filter and 50-60× concentrated using Amicon Ultra-15 100 kDa NMWCO columns (#UFC910024, Merck Millipore). The resulting concentrate was aliquoted and stored at -80°C until further use.

#### Cell transduction and puromycin selection

For each screen, 3 × 10^6^ MiaPaCa-2 cells in 1 ml of DMEM medium with polybrene (8 μg/ml) were seeded per well of a 6-well plate and appropriately diluted amount of concentrated lentivirus in 180 μl of DMEM media was added in order to obtain 0.3 MOI (the viral titer was assessed before the screen as well as during the screen). After 24 h incubation, medium with virus was discarded, cells counted, re-seeded either on 150 mm culture dishes (24 × 10^6^ cells) or 175 cm^2^ flasks (30 × 10^6^ cells), and cultivated under puromycin selection (1 μg/ml) for 3 days.

For initial screen with lentiCRISPRv2-GFPg1, 3 × 10^6^ cells were initially infected per sample (n = 1) and cultivated for 2 weeks after puromycin selection. In case of a screen with Brunello kinome library, 1.0 × 10^7^ cells (1000× coverage) were initially infected per sample (n = 3) and further cultivated for 2 weeks after puromycin selection.

#### Cell harvesting and genomic DNA isolation

For gDNA isolation either 3.0 × 10^7^ cells (lentiCRISPRv2-GFPg1 and Brunello kinome library) or 3.0 × 10^8^ cells (whole-genome Brunello library) per sample were harvested and processed according to Chen et al. ^26^. Briefly, cells were lysed in 50 mM Tris, 50 mM EDTA, 1% SDS, pH 8 in the presence of proteinase K (100 μg/ml) overnight at 55°C. Next day, samples were treated with RNAseA (50 μg/ml) for 30 minutes at 37°C and cooled on ice. After addition of 7.5M ammonium acetate, the clarified supernatant was further used for isopropanol precipitation of gDNA. Ethanol-washed gDNA pellets were dissolved in nuclease-free water and DNA concentration was assessed by Qubit.

### Library preparation

All primers and CRISPR-MIP probes used in this study are listed in Table S2.

#### Quantitative real-time PCR (qPCR)

The qPCR analysis was performed prior to any PCR amplification step during the library preparation in order to assess the proper cycle number. All qPCR reactions were performed using KAPA HiFi HotStart ReadyMix (Roche) with SYBR Green I (ThermoFisher Scientific). The reaction setup was identical to the conditions used for PCR1 or PCR2 from the actual CRISPR screen experiments.

#### Design of CRISPR-MIP probes

The design of CRISPR-MIP capture probes follows the general MIP structure^7–9^. The probe itself is a ssDNA molecule of 134 bp with extension (GTGGAAAGGACGAAACACC; Tm = 53.8°C) and ligation (AGCTAGGTCTTGAAAGGAGTGGG; Tm = 58.3°C) arms capturing the region of 112 bp, which includes complete sgRNA sequence. The Tm difference between probe arms should minimize the polymerase displacement of the ligation arm as discussed previously^8^. The backbone of the probe further encompasses 13 bp-long UMI, Illumina sequencing primers, and i7 index. P5/P7 Illumina adapter primers for PCR amplification step are localized within the captured sequence ensuring selective amplification of only properly elongated and ligated probes. The sequencing profile then counts with 60 and 15 bp for read1 and read2, respectively (Figure 1A).

#### CRISPR-MIP sgRNA abundance quantification

**Step 1 - denaturation & probe hybridization:** one reaction of 20 μl contains 1-10 μg of gDNA, 0.2 - 0.002 μM probe, and 1.5× ampligase buffer (Lucigen). The initial denaturation at 94°C for 5 minutes was followed by ramping down to 60°C (−0.1°C/s) at which the hybridization took place overnight.

**Step 2 - extension & ligation:** 10 μl of pre-heated (60°C) extension/ligation mix containing 5U of ampligase (Lucigen), 4U of Phusion HF DNA polymerase (#F530S, ThermoFisher Scientific), and 0.2 mM dNTPs was added to 20 μl of hybridization reaction. Properly mixed reaction was further incubated at 60°C for 1 hour.

**Step 3 - exonuclease treatment:** reaction was cooled down to 37°C and 10 U of Exonuclease I (# EN0581, ThermoFisher Scientific) and 50 U of Exonuclease III (#M0206L, NEB) was added to each reaction and incubated at 37°C for 45 minutes. This step removes non-circularized MIPs (Exo I) and genomic DNA (Exo III). Subsequently, both exonucleases were inactivated by incubation at 80°C for 20 minutes. The final reactions can be stored at -20°C until further use.

**Step 4 - PCR amplification:** the whole CRISPR-MIP reaction (31 μl) was then used as a template for 1 PCR reaction of 100 μl. The KAPA HiFi HotStart ReadyMix (Roche) was used with primers P5_tracrRNA_fwd and P7_tracrRNA_rev and the following cycling conditions: initial denaturation (95°C for 3 minutes), n cycles (98°C 20s; 60°C 15s; 72°C 30s), final extension (72°C 30 s). The number of cycles for the PCR amplification step was determined by qPCR for each screen independently. PCR cleanup was performed after the final PCR step by AMPure XP beads (Beckman Coulter). The concentration and purity of final PCR products was determined by Qubit and Bioanalyzer2100.

For the initial CRISPR-MIP screen with one-sgRNA model (lentiCRISPRv2-GFPg1) an equivalent of 0.15 × 10^6^ cells (1.0 μg) of either digested (*Bam*HI/*Hind*III) or undigested gDNA was used for 1 CRISPR-MIP reaction (i.e. 1 μg/reaction). In case of the Brunello kinome library screen, each gDNA sample started with 10.2 μg of gDNA (∼1.5 × 10^6^ cells) in three CRISPR-MIP reactions (3.4 μg/reaction), which were pooled (90 μl) and distributed into 3 PCR reactions. Following PCR amplification, these were then pooled back together.

#### PCR-based sgRNA abundance quantification

PCR-based amplification of sgRNA regions was performed as described previously^27^ with minor modifications - KAPA HiFi HotStart ReadyMix (Roche) was used for PCR1 and PCR2 with primers listed in Table S2 and following cycling conditions: PCR1 - initial denaturation (95°C for 3 minutes), n cycles (98°C 20s; 60°C 30s; 72°C 30s), final extension (72°C 30 s), PCR2 - initial denaturation (95°C for 3 minutes), n cycles (98°C 20s; 60°C 20s; 72°C 30s), final extension (72°C 30 s). The number of cycles for PCR1 and PCR2 was determined by qPCR for each screen independently. PCR cleanup was performed after PCR1 as well as PCR2 by AMPure XP beads (Beckman Coulter). The concentration and purity of final PCR products was determined by Qubit and Bioanalyzer2100.

In case of the Brunello kinome library screen, each gDNA sample started with 10.2 μg of gDNA (∼1.5 × 10^6^ cells) in three PCR1 reactions (3.4 μg/reaction), which were pooled prior the PCR1 cleanup. Subsequently, 10 μl from the respective PCR1 pool cleanup were added into PCR2, each reaction having a distinct P7-index primer.

### Counting of sgRNA reads

A custom Python script was written to extract sgRNA sequence, and UMI in case of CRISPR-MIP. For PCR, only R1 was analyzed, extracting the sgRNA as the 20bp in front of the sequence GTTTTAGAGC. For CRISPR-MIP, the sgRNA and UMI were extracted by position in R1 and R2. Only sgRNA sequences perfectly matching expected sgRNAs were retained (loss in the order of 1%). To perform efficient subsampling analysis, all reads were retained in memory, and then sampled without replacement five times for each depth. Deduplication was performed after read subsampling.

### UMI deduplication and modelling

Deduplication for CRISPR-MIP data was performed using UMI-tools^11^. Modeling was performed to find the best choice of algorithm parameters. Because the UMI is 13bp, we can model the distance from the UMI to a read with sequencing error as Binomal(n=13, p), where p is the sequencer error rate. Similarly, the distance from one UMI to another has a binomial distribution, Binomal(n=13, 3/4), based on the probability of each base being different from the reference UMI. This does not take sequencing errors into account, nor that all reads from another UMI are correlated. We thus extracted the most abundant sgRNA from a library to also qualitatively check these assumptions.

We qualitatively fitted the error rate p to be 3.5%. This is within range of Illumina sequencers, which generally have an error rate of 1% per base^28^. We used the more conservative fitted value to find cutoffs, as it would require more mismatches to be accepted.

A custom Python script was then used to simulate deduplication (simulate_UMI.ipynb), with n=3 bootstraps per setting. The average value is reported. To manage computational load, all sgRNAs are given the same expected count, as if all sgRNAs have the same coverage in the pool. For each setting (number of reads, number of UMIs, lower threshold, UMI-tools mismatch threshold), a defined number of molecules with a UMI are created following a Poisson distribution. Using the error rate p, these are then mutated, and the same number of reads are sampled for each UMI. Deduplication is then simulated: Each UMI is counted. Those with fewer than a defined threshold are removed. The remaining UMIs are deduplicated using the defined mismatch setting. The computed count DetectedUMI is then compared to the ground truth count #UMI as (DetectedUMI - #UMI) / #UMI.

### Saturation analysis

A custom python script was implemented for fast subsampling of the libraries. The entire FASTQ files were read into memory and sgRNA sequences extracted. Unknown sgRNAs were not included in the counts. For each library technical replicate, and each depth, 5 random subsamples were then extracted. MAGeCK was run on each bootstrap separately to enable computation of error bounds.

### GO analysis

The genes with FDR < 0.05 were compared to all genes present in the kinome library. The significant GO Biological Process annotations (GO:BP) were computed using DAVID^15,16^.

### Analysis of GC% bias

We searched for other CRISPR screens that might be GC-biased primarily using Pubmed and BioGRID ORCS^29^. The processed Sintov2022^17^ sgRNA counts were downloaded from GEO #GSM6008413. For the DepMap CRISPR screen^14^, raw counts were obtained from https://score.depmap.sanger.ac.uk/downloads, Release 1 (5th April 2019), raw_sgrnas_counts.zip.

The GC% bias was analyzed in R. Jitter has been added to the GC% in the plots, but models are fitted to the original values. For Sintov2022, the displayed linear model was generated using ggplot.

To correct for the GC% bias in the DepMap dataset, a model had to be fit of type foldChange ∼ f(GC%). Because of the asymmetry of the sgRNA log fold changes distribution, we fitted a skewed normal distribution using the R library sn^30^. Here we parameterized the mean by a 4th order polynomial over the GC% (this is the lowest polynomial which seemed to qualitatively fit the trend). For each sgRNA log fold change, we then subtracted this trend. We compute the per-gene average fold change as simply the fold change of the corresponding sgRNAs.

## Data availability

The raw sequencing data will be deposited on ArrayExpress. The pipeline for analyzing CRISPR-MIP sequencing data is available on GitHub, https://github.com/henriksson-lab/crispr-mip.

## Acknowledgements

All authors have read and agreed to the published version of the manuscript.

## Author contributions

M.S. did all work where otherwise not noted. I.Y. helped with follow-up experiments and finalization of the manuscript. I.N. helped with optimization. M.S. and J.H. wrote the manuscript and analyzed the data. J.H. supervised and conceived the project. All authors have read and agreed to the published version of the manuscript.

## References

1. Henriksson, J. et al. Genome-wide CRISPR Screens in T Helper Cells Reveal Pervasive Crosstalk between Activation and Differentiation. Cell 176, 882–896.e18 (2019).

2. Allen, F. et al. JACKS: joint analysis of CRISPR/Cas9 knockout screens. Genome Res. 29, 464–471 (2019).

3. Michlits, G. et al. CRISPR-UMI: single-cell lineage tracing of pooled CRISPR-Cas9 screens. Nat. Methods 14, 1191–1197 (2017).

4. Schmierer, B. et al. CRISPR/Cas9 screening using unique molecular identifiers. Mol. Syst. Biol. 13, 945 (2017).

5. Love, M. I., Huber, W. & Anders, S. Moderated estimation of fold change and dispersion for RNA-seq data with DESeq2. Genome Biol. 15, 550 (2014).

6. Svensson, V. Droplet scRNA-seq is not zero-inflated. Nat. Biotechnol. 38, 147–150 (2020).

7. Hardenbol, P. et al. Multiplexed genotyping with sequence-tagged molecular inversion probes. Nat. Biotechnol. 21, 673–678 (2003).

8. Akhras, M. S. et al. Connector inversion probe technology: a powerful one-primer multiplex DNA amplification system for numerous scientific applications. PLoS One 2, e915 (2007).

9. Li, J. B. et al. Multiplex padlock targeted sequencing reveals human hypermutable CpG variations. Genome Res. 19, 1606–1615 (2009).

10. Doench, J. G. et al. Optimized sgRNA design to maximize activity and minimize off-target effects of CRISPR-Cas9. Nat. Biotechnol. 34, 184–191 (2016).

11. Smith, T., Heger, A. & Sudbery, I. UMI-tools: modeling sequencing errors in Unique Molecular Identifiers to improve quantification accuracy. Genome Res. 27, 491–499 (2017).

12. Li, W. et al. MAGeCK enables robust identification of essential genes from genome-scale CRISPR/Cas9 knockout screens. Genome Biol. 15, 554 (2014).

13. Li, W. et al. Quality control, modeling, and visualization of CRISPR screens with MAGeCK-VISPR. Genome Biol. 16, 281 (2015).

14. Behan, F. M. et al. Prioritization of cancer therapeutic targets using CRISPR-Cas9 screens. Nature 568, 511–516 (2019).

15. Sherman, B. T. et al. DAVID: a web server for functional enrichment analysis and functional annotation of gene lists (2021 update). Nucleic Acids Res. 50, W216–W221 (2022).

16. Huang, D. W., Sherman, B. T. & Lempicki, R. A. Systematic and integrative analysis of large gene lists using DAVID bioinformatics resources. Nat. Protoc. 4, 44–57 (2009).

17. Sintov, E. et al. Whole-genome CRISPR screening identifies genetic manipulations to reduce immune rejection of stem cell-derived islets. Stem Cell Reports 17, 1976–1990 (2022).

18. Browne, P. D. et al. GC bias affects genomic and metagenomic reconstructions, underrepresenting GC-poor organisms. Gigascience 9, (2020).

19. Cui, Y., Xu, J., Cheng, M., Liao, X. & Peng, S. Review of CRISPR/Cas9 sgRNA Design Tools. Interdiscip. Sci. 10, 455–465 (2018).

20. Rabinowitz, R. & Offen, D. Single-Base Resolution: Increasing the Specificity of the CRISPR-Cas System in Gene Editing. Mol. Ther. 29, 937–948 (2021).

21. Koeppel, J. et al. Prediction of prime editing insertion efficiencies using sequence features and DNA repair determinants. Nat. Biotechnol. 41, 1446–1456 (2023).

22. Wu, K. & Malek, S. N. CRISPR/Cas9-Based Gene Dropout Screens. in Chronic Lymphocytic Leukemia: Methods and Protocols (ed. Malek, S. N.) 185–200 (Springer New York, New York, NY, 2019).

23. Walter, D. M. et al. Systematic In Vivo Inactivation of Chromatin-Regulating Enzymes Identifies Setd2 as a Potent Tumor Suppressor in Lung Adenocarcinoma. Cancer Res. 77, 1719–1729 (2017).

24. Doyle, T. et al. The interferon-inducible isoform of NCOA7 inhibits endosome-mediated viral entry. Nat Microbiol 3, 1369–1376 (2018).

25. Stewart, S. A. et al. Lentivirus-delivered stable gene silencing by RNAi in primary cells. RNA 9, 493–501 (2003).

26. Chen, S. et al. Genome-wide CRISPR screen in a mouse model of tumor growth and metastasis. Cell 160, 1246–1260 (2015).

27. Yau, E. H. & Rana, T. M. Next-Generation Sequencing of Genome-Wide CRISPR Screens. Methods Mol. Biol. 1712, 203–216 (2018).

28. Loman, N. J. et al. Performance comparison of benchtop high-throughput sequencing platforms. Nat. Biotechnol. 30, 434–439 (2012).

29. Oughtred, R. et al. The BioGRID database: A comprehensive biomedical resource of curated protein, genetic, and chemical interactions. Protein Sci. 30, 187–200 (2021).

30. Azzalini, A. A. The R package sn: The skew-normal and related distributions such as the skew-t and the SUN (version 2.1.1). Preprint at https://cran.r-project.org/package=sn (2023).

